# Reference genome assembly of a tetraploid accession of the tuber crop *Tropaeolum tuberosum*

**DOI:** 10.64898/2026.02.14.705015

**Authors:** Daniela Ruiz-Mateus, Ina Scheffler, María del Pilar Márquez-Cardona, Thomas Greb, Wilson Terán, Pascal Hunziker

**Author notes:** Author for correspondence:* Pascal Hunziker, Centre for Organismal Studies (COS), Heidelberg University, Im Neuenheimer Feld 360, 69120 Heidelberg, Germany. Authors contributed equally (Daniela Ruiz-Mateus, Ina Scheffler).

## Abstract

*Tropaeolum tuberosum* is a tuber-forming crop native to the Andes, valued for its nutritional content, pest resistance and adaptation to high-altitude environments. Despite its contribution to food security and sustainable agriculture, it remains an underutilized species with scarce genomic resources, limiting genetic and functional studies. To address this gap, we generated a reference genome assembly for a European *ex situ* tetraploid accession of *T. tuberosum* using PacBio HiFi sequencing. The assembly spans 1.3 Gb in 1,805 contigs (contigs N50 = 32.2 Mb, longest contig = 60 Mb) and recovers 87% of the estimated genome size. We assessed assembly completeness and accuracy using Benchmarking Universal Single-Copy Orthologs, which detected 98.5% complete genes (21.7% single-copy, 76.8% duplicated), 0.7% fragmented, and 0.7% missing, demonstrating near-complete gene space recovery consistent with a high-quality tetraploid genome. To evaluate transferability, we resequenced a field-collected Colombian genotype of economic relevance using Oxford Nanopore technology. Read mapping showed 99.7% of primary alignments to the reference (weighted mean coverage = 16.4×, 96.1% at ≥5×), confirming broad sequence conservation between accessions and validating the suitability of the *ex situ* reference as an anchor for *in situ* germplasm. Repetitive elements accounted for 71.3% of the genome. Using ANNEVO, we annotated 56,354 high-confidence protein-coding genes, achieving 98.3% complete BUSCOs, a PSAURON score of 97.2, and 90.5% taxonomic consistency with the rosid lineage (OMArk). This high-quality reference genome establishes a foundational resource for comparative genomics, population genomics and functional analyses on *T. tuberosum*, supporting future breeding and conservation of this important food resource.

**Significance statement:** *T. tuberosum* is an underutilized Andean tuber crop that combines highly nutritious tubers with remarkable disease resistance and UV tolerance, supporting Andean smallholder livelihoods under harsh high-altitude conditions. Beyond its role as a food crop, it is cultivated widely as an ornamental due to its abundant, brightly colored flowers closely resembling those of its relative nasturtium. Despite this dual relevance, genomic resources have been entirely lacking, limiting efforts to understand its domestication history, adaptation, and potential as a climate-resilient alternative to potato with higher nutraceutical properties. Our reference genome assembly provides the first genome-scale resource for *T. tuberosum*, creating a foundation for dissecting polyploid genome organization, key agronomic and nutritional traits, and interactions with biotic and abiotic stress. By enabling evolution, domestication, comparative genomics, breeding, conservation, and functional studies, this resource unlocks *T. tuberosum* as a model for resilient, diversified cropping systems globally.

## Introduction

*Tropaeolum tuberosum* Ruiz & Pav. (2n = 4x = 52), also known as *mashua, cubio* or *isaño*, is a tuber-forming crop traditionally cultivated by smallholder farmers in local communities in the Andes mountains. It thrives under harsh conditions – including strong diurnal temperature shifts and nutrient-poor soils – producing nutrient-rich tubers with high levels of carbohydrates, proteins, vitamins and diverse secondary metabolites with antioxidant, anti-inflammatory, diuretic and antimicrobial properties, including glucosinolates (Apaza Ticona et al. 2020; Luera-Quiñones et al. 2025; Coloma et al. 2022). These traits make *T. tuberosum* a promising species for food security as well as resilient and sustainable agriculture.

*T. tuberosum* belongs to the genus *Tropaeolum* (family Tropaeolaceae, order Brassicales), a Neotropical lineage of roughly 90 herbaceous species that includes ornamental nasturtiums and several tuber-forming taxa (Andersson & Andersson 2000). Its placement in the Brassicales is of particular biochemical interest: although glucosinolates are characteristic of this order, their diversity outside the Brassicaceae remains poorly characterized, highlighting the value of genomic data from non-brassicaceous taxa (Agerbirk & Koch 2025; Bird et al. 2025).

Like other edible Andean root and tuber crops, *T. tuberosum* exhibits remarkable morphological and genetic diversity (Pissard et al. 2008; Ruíz-Mateus et al. 2025; Fonseca & Márquez-Cardona 2024), yet low commercial value and limited market integration are jeopardizing its conservation as a food resource. In contrast to globally important tuber crops such as potato, its improvement has relied primarily on traditional farmer selection. Thus, despite its agronomic and nutritional potential, *T. tuberosum* remains largely underutilized (Luera-Quiñones et al. 2025).

No reference genome has been available for *T. tuberosum* and genomic resources remain scarce. Here we report a long-read–based, pseudo-haplotype–resolved genome assembly of a tetraploid accession of *T. tuberosum*. While a recent preprint described a genome for *Tropaeolum majus* (Friedhoff et al. 2024), this is the first assembly of *T. tuberosum*, providing a genomic framework for investigating specialized metabolism, tuber development, and environmental adaptation in a non-brassicaceous Brassicales lineage, while laying the foundation for genomics-assisted improvement of this neglected crop.

## Results and Discussion

We sequenced and assembled the genome of *Tropaeolum tuberosum* Ruiz & Pav (Tropaeolaceae, *Brassicales*) accession BGHEID007454 from the living collection of the Botanic Garden Heidelberg, Germany (**Figure 1a-b**). Using single-molecule real-time sequencing technology from Pacific Biosciences, a total of 128.2 Gb of PacBio HiFi sequencing data were produced. *K*-mer analysis revealed an estimated monoploid genome size of 418.1 Mb, consistent with an autotetraploid genome architecture and an estimated heterozygosity rate of 4.7% (**Figure 1c**). A genome survey based on *k*-mer frequency analysis of Illumina data estimated comparable metrics (**Supplementary Fig. S1**). *K*-mer-based genome size estimates (418-497 Mb) were moderately smaller than the monoploid genome size inferred from flow cytometry (2c = 2.48 Gb) (Veselý et al. 2013), which assuming tetraploidy yields a monoploid size of 620 Mb. Such underestimation is commonly observed in repeat-rich polyploid genomes due to compression of repetitive and near-identical homeologous regions. *K*-mer analysis of both HiFi and Illumina data supports an autotetraploid origin of *T. tuberosum*, with aaab forms (2.75%) exceeding aabb forms (1.72%) (**Figure 1c, Supplementary Fig. S1**), in line with the dominant cytological evidence (Johns & Neil Towers 1981).

**Fig. 1.**
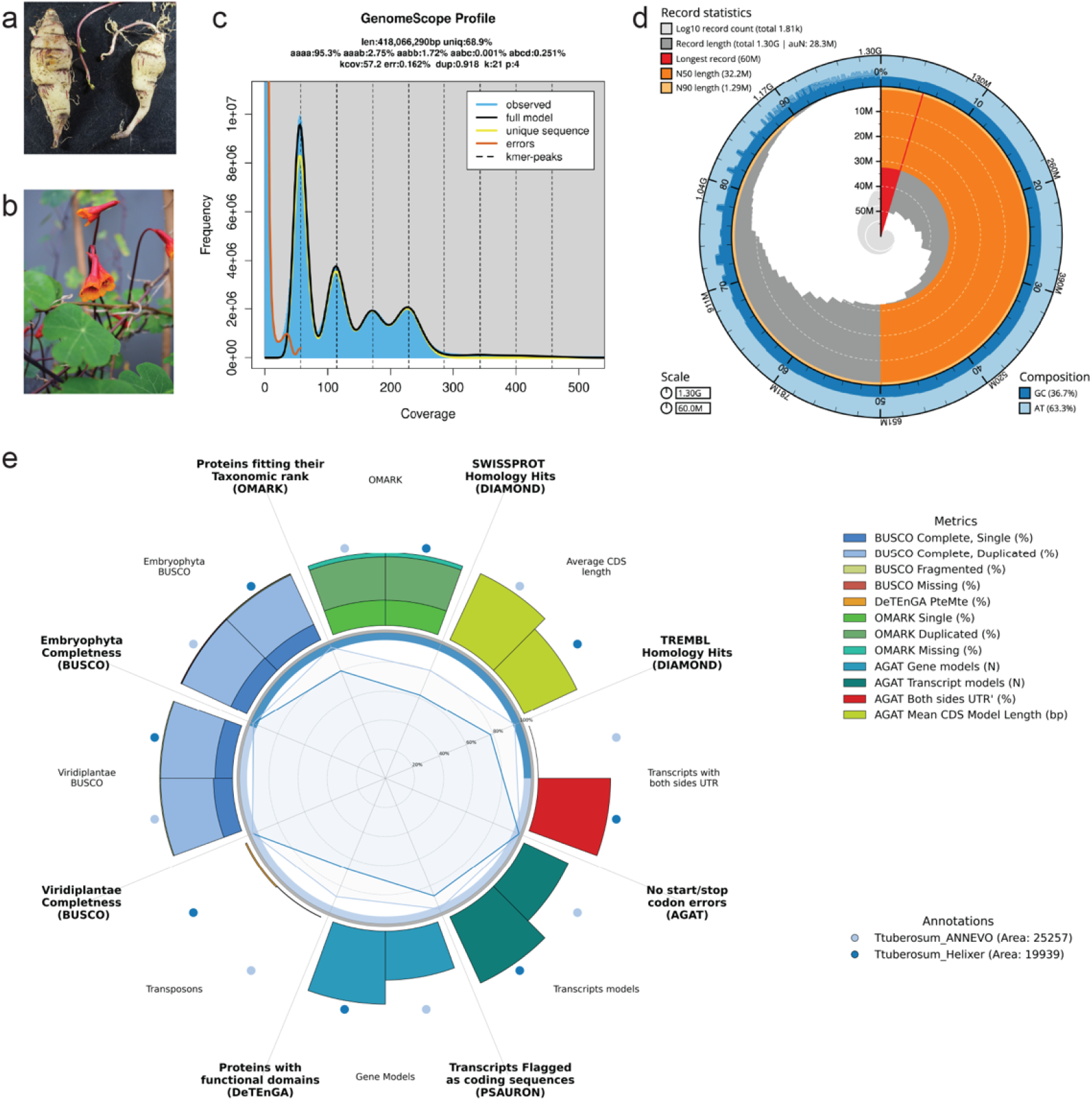
The *Tropaeolum tuberosum* reference genome. **(a)** Tubers of the BGHEID007454 accession. **(b)** Leaves, stems and flowers of the BGHEID007454 accession. **(c)** Distribution profile of 21-mer analysis. **(d)** Snail plot of the primary reference genome. **(e)** Benchmarking Helixer and ANNEVO annotations.

### Genome assembly

Initial genome assembly using hifiasm yielded a draft primary assembly containing 2,189 contigs and a pseudo-haplotype-resolved assembly comprising two haplotypes (hap1, 1.21 Gb and hap2, 0.98 Gb) of unequal size, as expected for autotetraploid genome assembly (**Supplementary Table S1-2**). Because the four homeologous copies do not segregate into two discrete sequence clusters, phasing algorithms partition them based on local heterozygosity patterns. Bandage plots showed numerous small contigs with low depth (**Supplementary Fig. S2**). Coverage distribution was bimodal, with a peak at 30–200× representing the nuclear genome and a tail of high-coverage contigs (>200×, n = 93) (**Supplementary Fig. S3**). 292 contigs showed very low coverage (<10×). Kraken2 classification confirmed that all 292 low-coverage contigs originated from non-target organisms (**Supplementary Table S3**). Contaminants (n = 292), plastid-derived (n = 60), mitochondria-derived (n = 30) and repeat contigs (n = 2) were removed (384 contigs in total, accounting for 17.5% of the original assembly by count).

The final primary assembly comprised 1,805 contigs (N50 of 32.2 Mb; largest = 60 Mb), with 90% of the assembly contained in 84 contigs (**Figure 1d, Table 1, Supplementary Table S1**). The size of 1.3 Gb matches the expected size of two monoploid complements (∼1.24 Gb (Veselý et al. 2013)), consistent with a collapsed pseudo-haploid assembly representing most sequence diversity across four homeologs. BUSCO analysis showed 98.5% completeness, 76.8% duplication, and 0.7% fragmentation, with identical results before and after decontamination, confirming no gene loss (**Table 1, Supplementary Table S1**). Merqury analysis yielded a consensus quality value (QV) of 60.4 and *k*-mer completeness of 75.4%

**Table 1.**
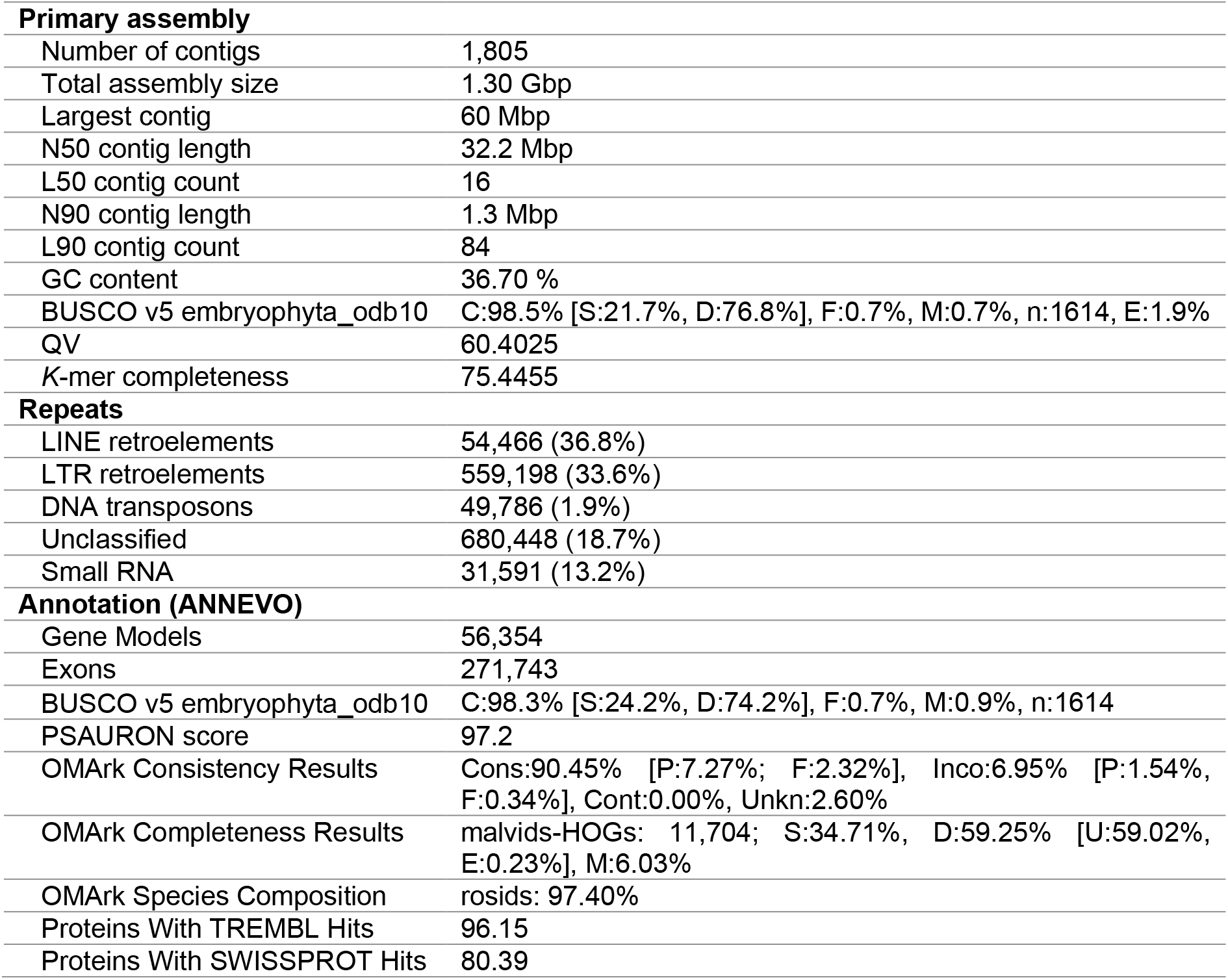
Statistics for the *Tropaeolum tuberosum* reference genome assembly and annotation.

(**Table 1, Supplementary Table S4**). The read-only and assembly copy number spectra aligned at ∼56× coverage, as expected for a collapsed tetraploid, indicating that reduced multiplicity reflects intended homeolog collapse rather than sequence loss (**Supplementary Fig. S4**). Assembly spectra exceeding read-only lines at higher multiplicities confirm comprehensive representation of multicopy sequences (**Supplementary Fig. S5**).

To assess the utility of the assembly as a reference for natural populations, Nanopore long reads (∼24×) from the field-collected genotype B15, originating from a farm in the department of Boyacá, Colombia, were mapped to the final primary assembly. A mapping rate of 99.7% and ≥98% of the assembly covered at a minimum depth of 5× demonstrate that the assembly captures the genomic content of geographically distant, *in situ* material with high fidelity, supporting its use as a reference for population-level and ecological genomic studies (**Supplementary Table S5**).

Of the 1,805 assembly contigs, 1,009 (55.9%) were covered at ≥10× mean depth, and a further 288 (16.0%) at 5–10× (**Supplementary Table S5**). The remaining 508 contigs (28.2%) showed low or absent coverage (<5×), collectively spanning only 23.5 Mb — 1.8% of the nuclear assembly by length. Repeat density in this fraction (998 masked bp/kb) was indistinguishable from that of well-covered contigs (952 bp/kb), against a genome-wide nuclear repeat content of 71.3%, indicating that the low-coverage fraction is not an enrichment of repetitive sequence but rather reflects the structural properties of short, repeat-dense contigs that are inherently difficult to anchor by read mapping regardless of genotype. Consistent with this, gene density on low-coverage contigs (164 models/Mb) was comparable to that of well-covered contigs (128 models/Mb), and only 3,446 of 56,354 annotated gene models (6.1%) resided in this fraction (**Supplementary Table S5**). The cross-genotype validation therefore confirms that the assembly provides comprehensive coverage of the genic and single-copy complement of the *T. tuberosum* genome, establishing it as a transferable reference resource for the species.

Together, the assembly statistics, BUSCO gene completeness, Merqury QV scores, *k*-mer completeness and cross-genotype validation demonstrate that the *T. tuberosum* genome assembly is highly accurate, complete and suitable for downstream analyses. Our primary assembly provides an annotation-grade reference genome sequence that was used for all gene prediction and functional annotation analyses described below.

The final haplotype-resolved assembly comprised hap1 (2,022 contigs, N50 = 18.4 Mb) and hap2 (580 contigs, N50 = 26.3 Mb), totalling 1.19 Gb and 0.96 Gb, respectively (**Supplementary Table S2**). Combined, the 2.19 Gb assembly represents ∼3.5× the monoploid genome size (620 Mb), recovering ∼88% of the expected tetraploid complement (∼2.48 Gb); the missing ∼12% likely corresponds to highly repetitive or heterochromatic regions. Gene completeness analysis showed slightly lower BUSCO scores per haplotype but 98.8% completeness when merged, indicating that unassembled sequence is largely non-genic (**Supplementary Table S2**). Merqury analysis revealed ∼70% *k*-mer recovery per haplotype and 95% combined, with consensus QV 60–65, reflecting high sequence accuracy. The *k*-mer spectrum displayed four coverage peaks (∼56×, ∼112×, ∼168×, ∼224×), matching the expected 1×–4× pattern for an autotetraploid at ∼56× per haploid copy (**Supplementary Fig. S6**). Minimal read-only fractions confirmed near-complete representation of the read *k*-mer space, while balanced hap1- and hap2-specific fractions and a dominant shared component at higher multiplicities (**Supplementary Fig. S7**) are characteristic of autopolyploidy. Distinct haplotype-specific peaks verified successful phasing, and per-haplotype spectra showed that missing *k*-mers reflect sequence assigned to the alternate haplotype rather than gaps (**Supplementary Figs. S8–S9**). Together, these results indicate a high-quality pseudo-haplotype-resolved, partially phased assembly of an autotetraploid genome

### De novo repeat and gene annotation

Using RepeatModeler, we identified 1,706 repeat families, including 732 structurally defined LTR retrotransposons (**Table 1, Supplementary Table S6**). Repetitive elements accounted for 71.3% of the genome, dominated by Gypsy and Copia LTR retrotransposons. This composition aligns with other eudicot genomes of similar size (Chanderbali et al. 2022), where LTR retrotransposons contribute substantially to genome expansion (Zhang et al. 2020). The final nonredundant repeat library was used for genome-wide masking and annotation.

Helixer and ANNEVO predicted 84,059 and 56,354 protein-coding genes, respectively (**Figure 1e, Table 1, Supplementary Table S7**). Both annotations showed near-complete gene space coverage, with BUSCO completeness scores of 98.1% and 98.3%, respectively. In both cases, the majority of complete BUSCOs were duplicated (∼75%), consistent with the retention of duplicated genes following ancient whole genome duplication events common in flowering plants (Van de Peer et al. 2017). Despite comparable completeness, ANNEVO consistently produced higher-quality gene models across all metrics assessed. ANNEVO achieved a substantially higher ORF accuracy score (PSAURON: 97.2 vs. 87.5), greater taxonomic consistency with the rosid lineage (OMArk: 90.5% vs. 71.5%), and a higher proportion of proteins with significant homology hits in both TrEMBL (96.2% vs. 78.2%) and Swiss-Prot (80.4% vs. 62.0%). Helixer’s higher gene count was accompanied by clear signs of inflation: 4,557 overlapping gene models (vs. 59 in ANNEVO), 504 models with missing stop codons, 4.2% of proteins shorter than 50 amino acids (vs. 0.1%), and nearly four times more transposable element-derived transcripts (DeTEnGA PteMte: 1.02% vs. 0.27%). The higher proportion of unknown hits in Helixer’s OMArk analysis (19.95% vs. 2.60%) further suggests that a substantial fraction of its additional gene models lack biological support.

Based on these results, the ANNEVO annotation was selected as the primary gene set. The 56,354 high-confidence protein-coding genes predicted by ANNEVO are consistent with the tetraploid genome of *T. tuberosum*. The closely related diploid *Tropaeolum majus* harbors 32,924 protein-coding genes (Friedhoff et al. 2024). Assuming a similar ancestral gene complement, whole genome duplication would be expected to approximately double this number, with subsequent gene loss and fractionation yielding the observed count in *T. tuberosum*.

### Assembly and annotation of organelle genomes

The plastid genome was assembled as two complete circular paths of identical length (∼150 kb), representing the two orientations of the large single-copy / inverted repeat / small single-copy (LSC/IR/SSC) quadripartite structure characteristic of land plant chloroplasts (**Supplementary Figure S10**). This result is consistent with the well-documented flip-flop recombination of the IR region and confirms assembly completeness.

Mitochondrial genome assembly recovered one primary scaffold of 550,896 bp and a complex assembly graph with over 100 repeat patterns (**Supplementary Figure S11**). Such structural complexity is typical of plant mitochondria, which undergo repeat-mediated recombination and exist as dynamic genomic populations. Thirty high-coverage HiFi contigs mapped to this scaffold, confirming their mitochondrial origin and warranting their exclusion from the nuclear assembly. The most extreme outlier (ptg000094l, 1,293×) showed 33 BLASTn hits to the mitochondrial genome, indicating a highly reiterated region with elevated read depth.

## Conclusions

This first reference genome assembly for *T. tuberosum* achieves high completeness and sequence quality, providing a foundation for comparative genomics, functional analyses and diversity studies. This resource enables investigation of polyploid genome organization, tuber development, and specialized metabolism in *T. tuberosum* and comparative studies across the Tropaeolaceae family.

## Materials and Methods

DNA in vitro-grown *T. tuberosum* accession BGHEID007454 (Botanic Garden Heidelberg, Germany) was extracted using a CTAB-based protocol (Vaillancourt & Buell 2019). The Colombian B15 morphotype was collected from small producers’ farms in Boyacá, Colombia (Fonseca & Márquez-Cardona 2024), and maintained in a conservation plot at San Javier Farm, Cogua, Cundinamarca, Colombia (05°03’37.0” N, 073°56’41.6” W; 2590 m). DNA of B15 was extracted from fresh leaf tissue as described (Kalendar et al. 2021).

BGHEID007454 DNA was sequenced on an Illumina NextSeq2000 at the Deep Sequencing Core Facility, Heidelberg University. Long-read sequencing used the SMRTbell Express Template Prep Kit 2.0 on a PacBio Revio by Novogene (https://www.novogene.com/). B15 DNA libraries (SQK-LSK114) were sequenced on a PromethION 24 (V14 chemistry, R10 flow cells) by SNP Saurus (http://www.snpsaurus.com). Genome size, heterozygosity, and duplication rate were estimated using jellyfish and GenomeScope2.

HiFi reads were assembled with hifiasm. The primary assembly underwent contaminant (Kraken2; n=292) and organelle (BLASTn; n=89) removal. Quality metrics (gfastats, BUSCO, merqury, BlobToolKit) and B15 ONT mapping (minimap2; mosdepth) are detailed in the Supplementary Data.

The assembly was soft-masked (RepeatModeler, RepeatMasker). Gene models were predicted ab initio using Helixer and ANNEVO. Annotation quality was benchmarked with GAQET2 including BUSCO, OMArk, PSAURON, DeTEnGA/TEsorter, and DIAMOND (see Supplementary Data).

Plastid and mitochondrial genomes were assembled from Illumina reads using GetOrganelle. The plastid was annotated with GeSeq. Full details are in the Supplementary Data.

## Data Availability

Raw reads for the BGHEID007454 and B15 genotypes used in this study have been deposited in the European Nucleotide Archive (ENA) at EMBL-EBI under accession numbers PRJEB108055 (https://www.ebi.ac.uk/ena/browser/view/PRJEB108055) and PRJEB109811 (https://www.ebi.ac.uk/ena/browser/view/PRJEB109811), respectively. Reference genome assembly (v1.2), haplotype sequences and annotation files have been deposited at FigShare (https://doi.org/10.6084/m9.figshare.31338463).

### Ethics and Compliance

The *Tropaeolum tuberosum* accession BGHEID007454 was obtained under a Material Transfer Agreement from the Botanic Garden Heidelberg, Germany. Maintained in European collections since at least 1995 (predating the Nagoya Protocol and EU Regulation 511/2014), it lacks documented collecting event or provenance. The collection and access to the Colombian genetic resource *Tropaeolum tuberosum* B15 were covered under the Access to Genetic Resources and Derived Products Agreement No. 256, “Studies on the Genetic Diversity of Native Crops of Colombia,” and its Addendum No. 1, signed between the Ministry of Environment and Sustainable Development of Colombia and Pontificia Universidad Javeriana.

## Author contributions

Study conception and design: PH. Sample preparation and experimental work: DRM, IS, MPM, PH. Data analysis: DRM, WT, PH. Manuscript draft: PH. Manuscript final revision and consent: DRM, IS, MPM, TG, WT, PH.

## Acknowledgements

We thank the Botanical Garden Heidelberg for plant material (BG HEID 007454), Sandra Schmöckel and David Jarvis for helpful discussions, and Aureliano Bombarely for early ANNEVO testing. We thank David Ibberson (Deep Sequencing Core Facility, Heidelberg University) for technical support and Marcus Koch for commenting on the manuscript. We acknowledge the SDS@hd data storage service supported by MWK and DFG (INST 35/1803-1 FUGG, INST 35/1804-1 LAGG) and computational resources provided by the High Performance and Cloud Computing Group, University of Tübingen, through bwHPC and DFG (Project 455787709, bwForCluster BinAC2). This work was funded by the Klaus Tschira Boost Fund (Klaus Tschira Stiftung/GSO e.V.) awarded to PH, and Fontagro (IDB) under grant “Root to Food: Improvement of Yield in Potatoes and Other Andean Tubers” (ATN/RF-18120-RG) to MPMC and WT.

