## Supplemental Data for "Reference genome assembly of a tetraploid accession of the tuber crop *Tropaeolum tuberosum*"

[†]Authors contributed equally

### Supplementary Materials and Methods

#### Plant growth

20 cm long shoot tip segments were cut from a vegetative *T. tuberosum* BG HEID 007454 plant, leaves removed with scissors, and sterilized by sequentially soaking 5 min in 70 % ethanol and 15 min in 1.5 % sodium hypochlorite, 0.1 % Tween-20, followed by six washes with sterile water. Nodal segments of 1-2 cm length were cut and placed onto 1X MS medium (Musharige and Skoog medium including vitamins (Duchefa M0222), 0.5 g/L MES, 30 g/L sucrose, 8 g/L agar, pH 5.7) in Magenta GA-7 vessels. Plants were grown in a climate chamber at 25°C under long-day conditions (16 h light). 3-week-old plants were used for DNA extraction.

The Colombian morphotype (B15) was collected by Fonseca and Márquez-Cardona (2024) from small producers' farms in the department of Boyacá, Colombia. B15 tubers were subsequently planted and maintained in a conservation plot at San Javier Farm, Cogua (05°03'37.0" N, 073°56'41.6" W; 2590 m), in the department of Cundinamarca, Colombia.

#### DNA extraction, Illumina, PacBio and Nanopore sequencing

We extracted high molecular weight DNA from *T. tuberosum* accession BG HEID 007454 maintained in the living collection of the Botanic Garden Heidelberg, Germany. The accession was received from the Botanic Garden Dresden, Germany, in 2025, which had received it from the Botanic Garden Bonn, Germany, in 1995. The accession has been maintained under vegetative propagation in European botanical collections for at least three decades. Historical records prior to 1995 and original geographic provenance are unavailable. High molecular weight DNA was extracted from a single in vitro grown plant using a CTAB-based protocol (Vaillancourt & Buell 2019). DNA quality and quantity were assessed using a NanoDrop spectrophotometer (ND-2200, Thermo Fisher Scientific), Qubit dsDNA BR Assay (Thermo Fisher Scientific), and TapeStation Genomic DNA ScreenTape Assay (Agilent Technologies). For short-read sequencing, 117 ng DNA in 15 µl was processed using the NEBNext FS DNA Express reagents with NEBNext UDI Oligos for Illumina. 20 min of fragmentation was used and 6 cycles of PCR. Libraries were purified using Macery-Nagel Magbeads. The library was quantified using the ThermoFisher Qubit HS DNA Assay and quality checked with the D1000 Assay on the Agilent Tapestation. 650 pM were loaded onto the Illumina NextSeq 2000 using a P1 300 cycle flowcell in 150 PE mode. For long-read sequencing, DNA was shipped to Novogene Corporation (https://www.novogene.com/). DNA quality was assessed a second time by Novogene Corporation using NanoDrop, Qubit, gel electrophoresis and pulsed-field gel electrophoresis. Library preparation was performed using the SMRTbell Express Template Prep Kit 2.0. and the resulting library was sequenced using one SMRT Cell 25M Tray on a PacBio Revio system.

For Nanopore sequencing, fresh leaves of B15 were collected, ground in liquid nitrogen and stored at −80 °C in the Plant Molecular Biology Laboratory at Pontificia Universidad Javeriana, Bogotá, Colombia, for subsequent analyses. DNA extraction was performed from leaf tissue of a single individual as previously described (Kalendar et al. 2021). DNA quality, quantitation and integrity were assessed by spectrophotometry (Nanodrop ND-1000, Thermo Fisher Scientific) and agarose gel electrophoresis. Samples showing high integrity and an A260/A280 ratio ≥ 1.8 were considered suitable for sequencing and shipped to SNP Saurus DNA sequencing facility (http://www.snpsaurus.com). DNA quantitation and quality was further assessed using TapeStation Genomic DNA ScreenTape Assay (Agilent Technologies) and library preparation was carried out using the SQK-LSK114 Ligation Sequencing kit (Oxford Nanopore Technologies). Whole-genome sequencing (~20× coverage) was conducted using the Oxford Nanopore Technologies PromethION 24 platform with V14 chemistry and R10 flow cells, employing super-accuracy basecalling. Raw sequencing data were evaluated for quality using Nanoplot v1.46.1 (De Coster & Rademakers 2023). Adapter sequences were removed using Porechop v0.2.4 (Wick et al. 2017). Reads shorter than 1000 bp were removed using Filtlong v0.3 (Wick 2018), and Kraken2 v2.1.6 (Wood et al. 2019) was used to remove sequences originating from off-target organisms. A custom database containing bacterial, archaeal, fungal, and human RefSeq genomes was used for classification, applying a confidence threshold of 0.5.

#### Genome size estimation

Illumina short reads were checked using fastqc v0.12.1 [14] and trimmed using fastp v0.22.0 using a qualified phred quality of Q20 and required length of 50 bp [15]. Jellyfish v2.2.10 was used to perform *k*-mer = 21 frequency distribution analysis (Marçais & Kingsford 2011). Subsequently, GenomeScope v2.0 was used to estimate genome size, heterozygosity, and duplication rate using k = 21 and n = 4 (Ranallo-Benavidez et al. 2020).

#### Genome sequence assembly

The genome was assembled using hifiasm v0.25.0-r726 (Cheng et al. 2024, 2022, 2021) with hifiasm -o asm -t 48 --hg-size 0.42g. Additional purging strategies and assembly parameter tuning (including --hom-cov and -s adjustments) were evaluated. However, coverage-based diagnostics, including the absence of a bimodal read-depth distribution, did not support reliable discrimination between allelic and redundant sequence. Hifiasm was run with default settings (-l 2), incorporating two internal haplotig purging rounds. Aggressive external filtering with purge_dups v1.2.6 (Guan et al. 2020) resulted in severe over-collapsing (~91 Mb), and was therefore not applied.

Assembly statistics were generated using gfastats v1.3.11 (Formenti et al. 2022). Genome quality was assessed using BUSCO v5.8.3 (Manni et al. 2021) with the embryophyta_odb10 database in genome mode. *K*-mer-based evaluation of genome completeness and accuracy was performed using meryl v1.4.1 and merqury v1.3 (Rhie et al. 2020). Meryl was used to count canonical *k*-mers (k = 21) from the raw HiFi reads and from the assembled haplotype sequences. Merqury then compared the read k-mer spectra with each haplotype assembly to estimate the proportion of read *k*-mers captured, the completeness of each haplotype, and base-level consensus quality (QV).

Sequencing depth per contig was assessed by mapping HiFi reads to the primary assembly using minimap2 v2.30 (Li 2018) and computing per-contig coverage statistics with samtools v1.23 (Li et al. 2009). Contigs were flagged as putative contaminants (mean depth < 10×, n = 292) or putative organelle-derived (mean depth > 200×, n = 93) based on coverage distribution relative to the overall assembly mean (~59×).

Taxonomic classification of flagged contigs was performed using Kraken2 v2.1.6 (Wood et al. 2019) against the standard database (RefSeq complete genomes for bacteria, archaea, viruses, human, and UniVec_Core; compiled 2025-09-24) and the fungi database. All 292 low-coverage contigs received taxonomic assignments and were removed from the assembly.

High-coverage contigs (n = 93) were compared against the assembled organelle genomes (see below) using BLASTn v2.17.0 (Chen et al. 2015) (≥ 80% identity, ≥ 30% query coverage for mitochondria; ≥ 90% identity, ≥ 50% query coverage for plastid). Fifty-nine contigs were classified as plastid-derived and 30 as mitochondria-derived; an additional plastid contig (ptg000098l) was identified by remote BLASTn against the NCBI nt database matching a congruous *Tropaeolum majus* chloroplast genome (GenBank ON641313.1). Two remaining high-coverage contigs with no informative hits (ptg000532l, ptg000614l) were classified as putative ribosomal DNA repeats based on their coverage profile and absence of specific taxonomic signal, and were excluded. A total of 384 contigs were removed from the primary assembly, yielding a final nuclear assembly of 1,805 contigs.

Assembly quality was assessed using BlobToolKit v4.5.0 (Challis et al. 2020). A BlobDir dataset was created directly from the filtered assembly FASTA file, and a snail plot was subsequently generated from the BlobDir.

#### Cross-genotype validation by ONT read mapping

ONT reads were mapped to the reference genome assembly using minimap2 v2.30 (Li 2018) with the ont-map preset. Mapping statistics were obtained with samtools v v1.23 (Li et al. 2009) flagstat. Per-contig mean coverage was computed using mosdepth v0.3.13 (Pedersen & Quinlan 2018) from the coordinate-sorted alignment. The complete contig list (1,805 entries) was derived from the assembly FASTA index to retain contigs absent from the mosdepth summary due to zero coverage. Contigs were classified by mean ONT depth into four categories: high-copy (>200×, n = 27), well-covered (≥10×, n = 982), partial coverage (5–10×, n = 288), and low/absent coverage (<5×, n = 508). The 27 high-copy contigs were excluded from repeat and gene density analyses to avoid confounding these statistics with non-nuclear sequence composition; they are included in all coverage summaries and counted within the well-covered (≥10×) fraction.

Overlapping repeat intervals were merged with bedtools v2.31.1 merge (Quinlan & Hall 2010) prior to all coverage calculations to prevent double-counting of nested transposable elements. Per-contig repeat density (masked bases per kilobase) and length-weighted repeat fraction were computed with bedtools coverage. Gene model coordinates were derived from the ANNEVO annotation; mRNA features (n = 56,354) were used as the counting unit, representing one predicted locus per model. Gene density (models per Mb) and the number of gene models per coverage class were computed by intersecting gene and contig coordinates with bedtools intersect. All repeat and gene analyses were performed on the nuclear contig set (1,778 contigs, 1,270 Mb) after exclusion of high-copy contigs.

#### Repeat modelling, masking and genome annotation

The filtered primary assembly was utilized for *de novo* repeat modeling using RepeatModeler v2.0.7 (Flynn et al. 2020) using rmblast 2.14.1+. Assembly was soft-masked using RepeatMasker v4.2.2 (Tarailo-Graovac & Chen 2009). Gene prediction was performed independently using two deep learning-based ab initio annotation tools: Helixer v0.3.2 (Holst et al. 2025) and ANNEVO v2.2.1 (Ye et al. 2025).

For Helixer, the soft masked assembly was processed using the pre-trained 'land_plant' model with default parameters optimized for plant gene structures. Helixer was run without overlap mode (--no-overlap) using a batch size of 4 on an NVIDIA A100 80GB PCIe GPU with CUDA 11.8 and cuDNN 8.9.7. Gene models were output in GFF3 format, and protein sequences were extracted using gffread v0.12.7 (Pertea & Pertea 2020). For ANNEVO, the soft-masked assembly was processed using the pre-trained 'Embryophyta' model on an NVIDIA A100 80GB PCIe GPU with CUDA 12.6 and PyTorch v2.1.0. Gene models were output in GFF format.

The quality of both annotations was assessed using GAQET2 v2.0 (Garcia-Carpintero et al. 2025), which integrates multiple complementary benchmarking tools. Gene model structural integrity was evaluated using AGAT v1.4.1 (Dainat 2026). Proteome completeness was assessed with BUSCO v5.8.3 (Manni et al. 2021) using both the embryophyta_odb10 (n=1,614) and viridiplantae_odb10 (n=425) lineage databases in protein mode. Taxonomic consistency and completeness were evaluated using OMArk v0.3.1 (Nevers et al. 2025) with

LUCA.h5 OMA database against the malvids hierarchical orthologous groups (HOGs; n=11,704), with a target taxon ID of 147035 (*Tropaeolum tuberosum*). Open reading frame accuracy was scored using PSAURON v1.0.4 (Sommer et al. 2025), and transposable element contamination among predicted gene models was assessed using DeTEnGA v1.0 with the rexdb-plant database via TEsorter v1.4.7 (Zhang et al. 2022) and InterProScan v577-108.0 (Jones et al. 2014). Protein homology was assessed by sequence similarity searches against the UniProt Swiss-Prot and TrEMBL databases using DIAMOND v2.1.11 (Buchfink et al. 2021).

### Organelle genome assembly and annotation

Plastid and mitochondrial genomes were assembled de novo from Illumina paired-end reads using GetOrganelle v1.7.7.1 (Jin et al. 2020) with the embplant_pt and embplant_mt seed and label databases (version 0.0.1). For plastid assembly, two complete circular paths were recovered corresponding to the two orientations of the inverted repeat region; path 1 was selected as the representative sequence and annotated with GeSeq (https://chlorobox.mpimp-golm.mpg.de/geseq.html). Mitochondrial assembly was performed with up to 100 extension rounds and k-mer sizes of 21, 45, 65, 85, and 105 bp. Read input was capped at 50× estimated organelle coverage (--reduce-reads-for-coverage 50, --max-reads 20,000,000) to reduce memory requirements. The resulting assembly graph was visualised in BandageNG v2025.6.1 (Wick et al. 2015).

### Supplementary Figures

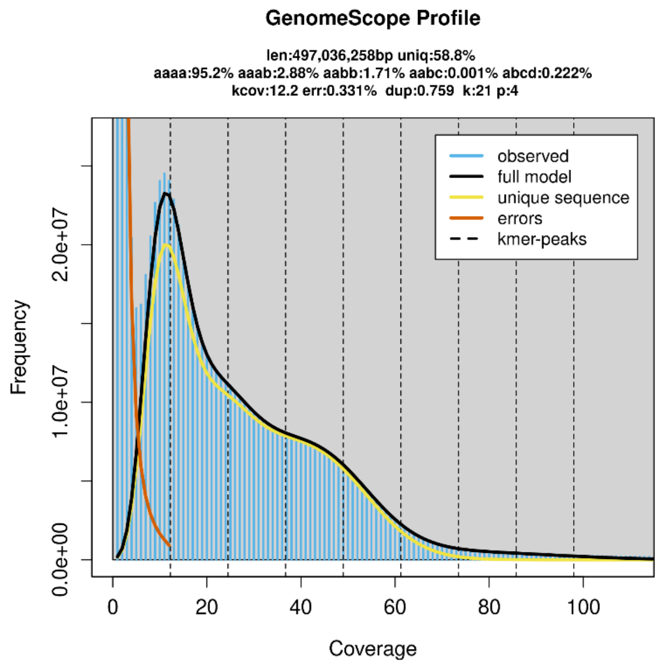


**Supplementary Figure S1:** Genome survey based on *k*-mer frequency analysis of 36.6 Gb of filtered Illumina short-read data.

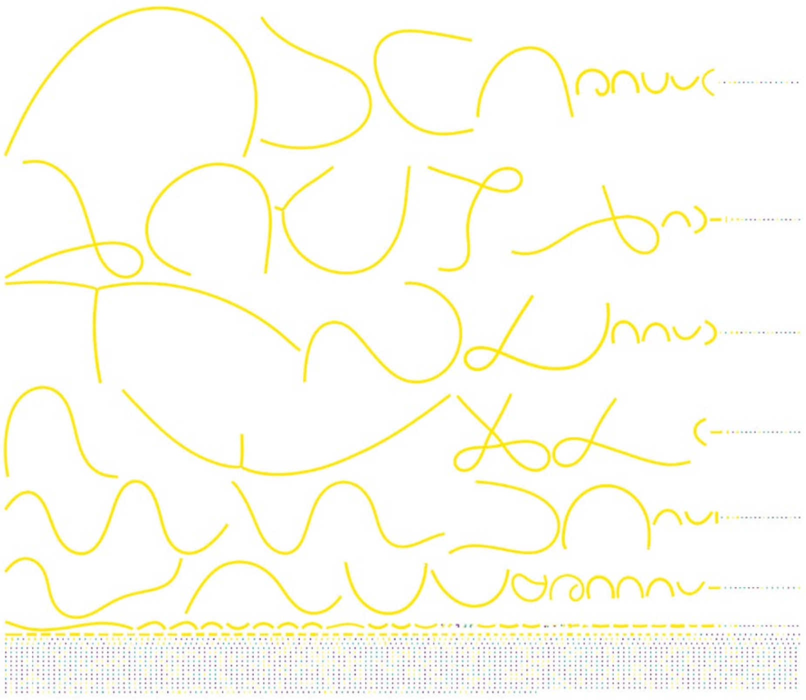

**Supplementary Figure S2:** Bandage plot colored according to depth using default settings. Yellow color shows nodes with high depth above the 3rd quartile. Median depth is 53x.

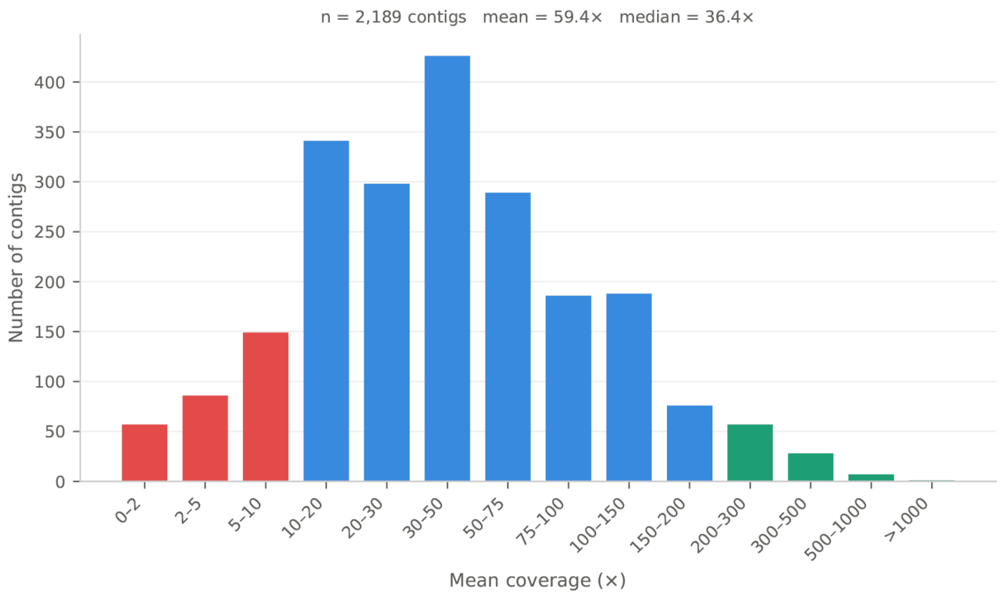


**Supplementary Figure S3:** Coverage distribution of 2,189 primary contigs. Bars show contig counts per coverage bin.

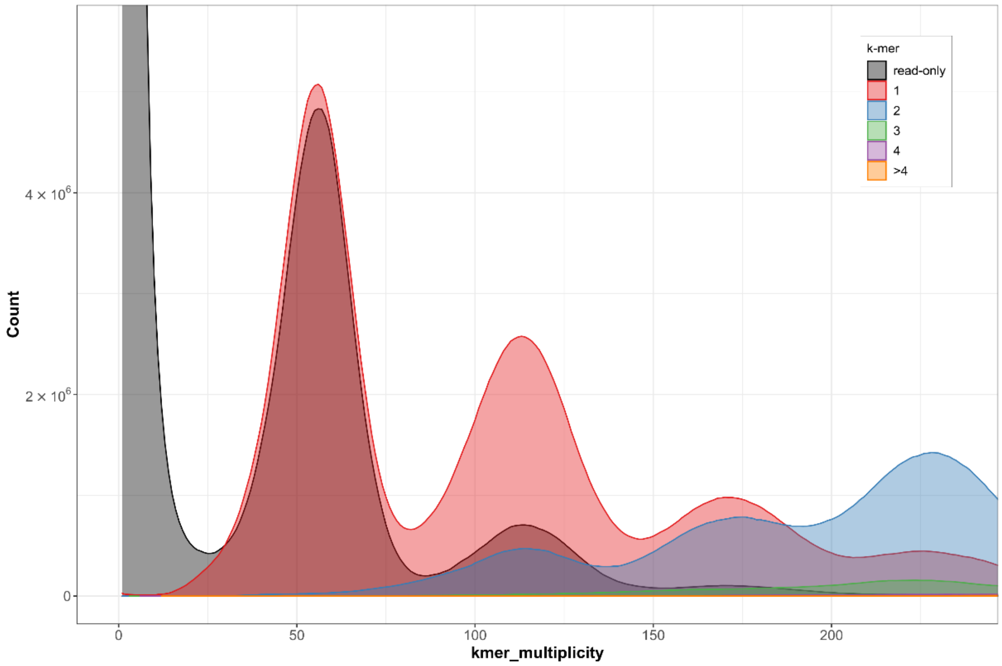


**Supplementary Figure S4:** *K*-mer copy-number spectra of the primary assembly. The overlap of the read-only line with the 1-copy assembly fraction at ~56× and ~112× reflects the expected collapse of homeologous copies in a pseudohaploid reference assembly.

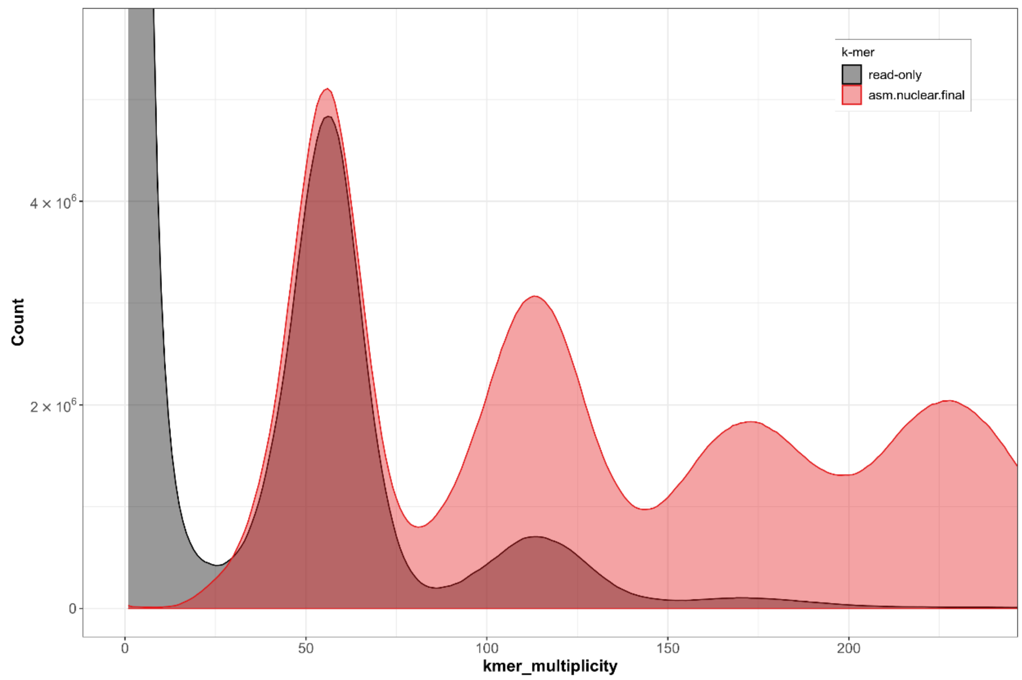


**Supplementary Figure S5:** Assembly *k*-mer spectra of the primary assembly. The assembly *k*-mer line (red) substantially exceeds the read-only line (black) at all higher-multiplicity peaks, confirming that multi-copy genomic content is well-represented in the collapsed primary assembly.

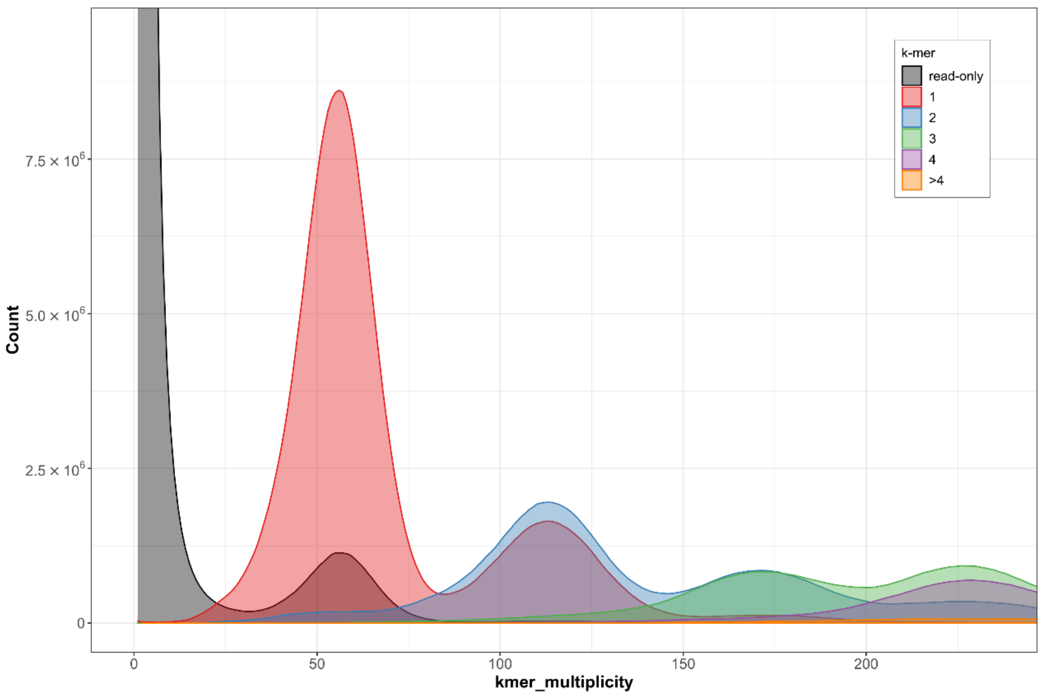


**Supplementary Figure S6:** *K*-mer copy-number spectra of the haplotype-resolved assembly. *K*-mer multiplicity (x-axis) is plotted against *k*-mer count (y-axis). Coloured regions indicate *k*-mers assigned to 1–4 copies in the assembly; grey indicates read-only k-mers (present in reads but absent from the assembly). Four peaks corresponding to the expected 1×–4× coverage intervals (~56×, ~112×, ~168×, ~224×) confirm the autotetraploid ploidy level. The low-multiplicity read-only spike reflects sequencing error *k*-mers.

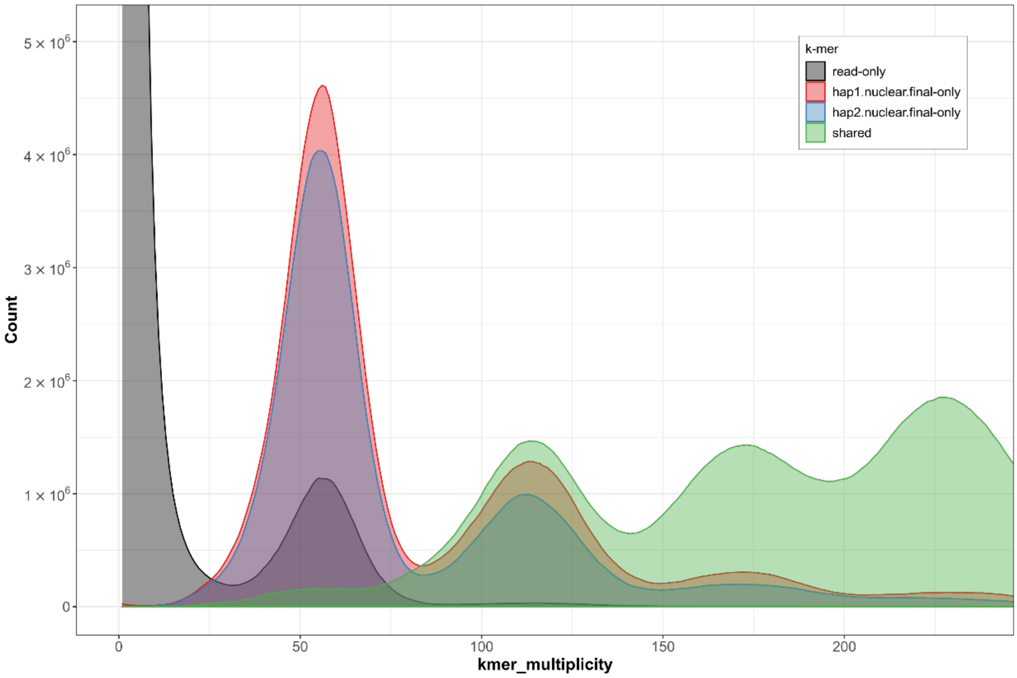


**Supplementary Figure S7:** Assembly *k*-mer spectra of the haplotype-resolved assembly. *K*-mers are coloured by haplotype assignment: hap1-only (red), hap2-only (blue), and shared between both haplotypes (green). Read-only *k*-mers are shown in black. The large shared fraction at higher multiplicities reflects the high homeologous sequence similarity characteristic of autopolyploids.

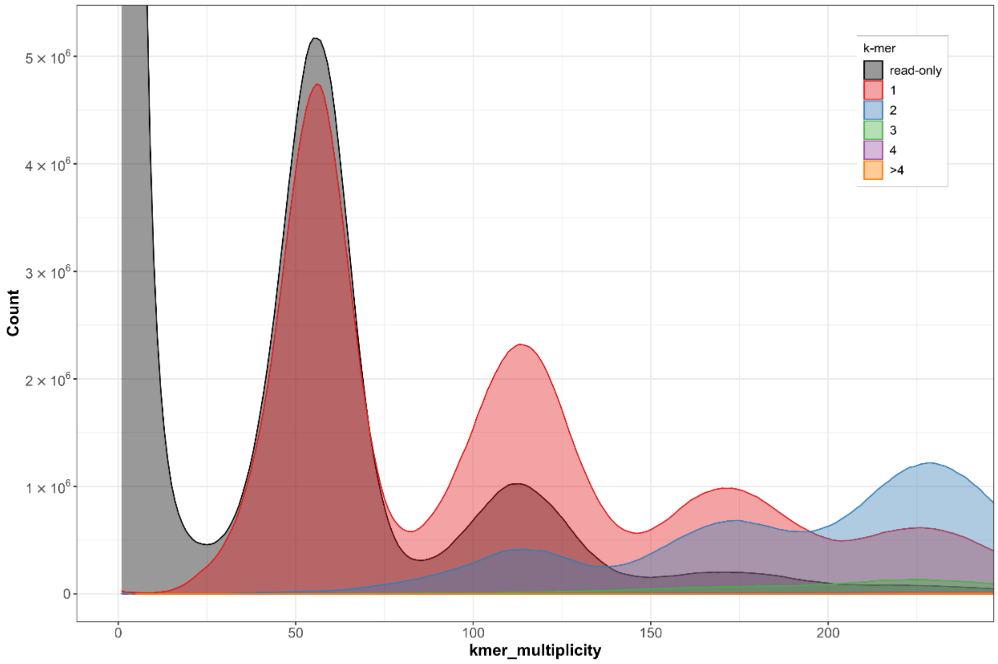


**Supplementary Figure S8:** *K*-mer copy-number spectra of haplotype 1 evaluated against all HiFi reads. The read-only fraction at the primary coverage peak corresponds to sequence correctly phased into haplotype 2.

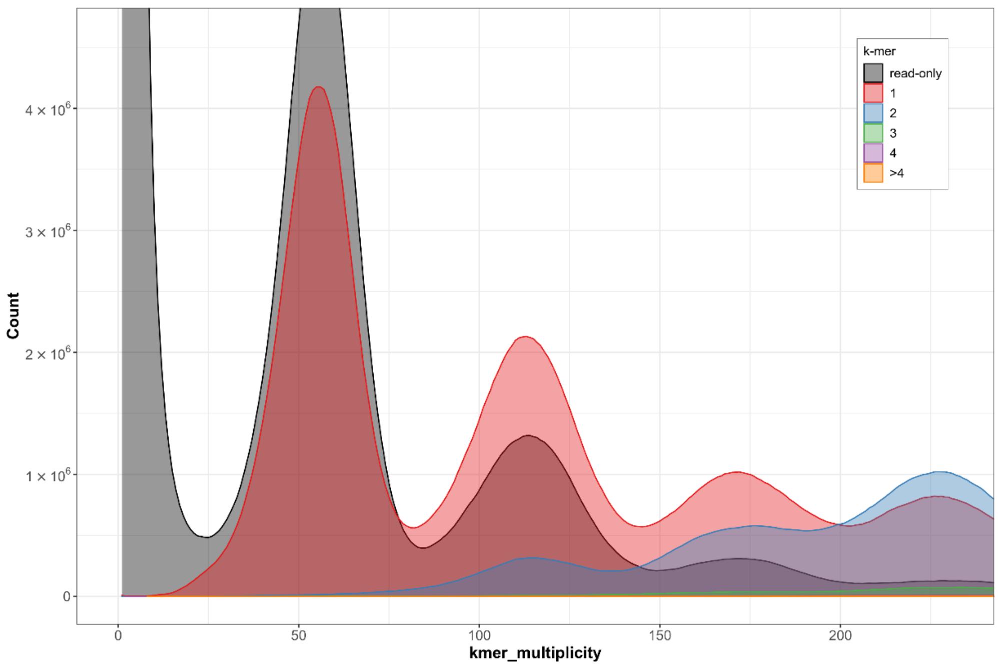


**Supplementary Figure S9:** *K*-mer copy-number spectra of haplotype 2 evaluated against all HiFi reads. As for haplotype 1, the read-only fraction reflects sequence distributed to the alternate haplotype.

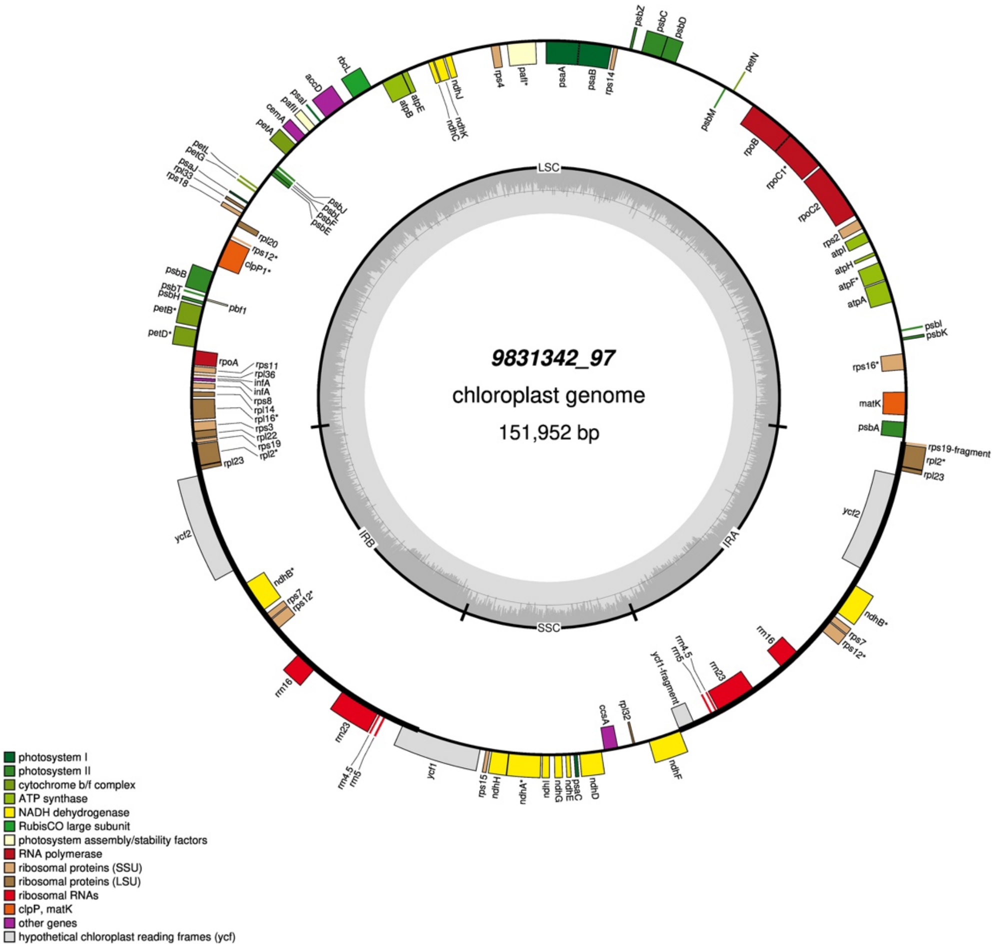


**Supplementary Figure S10:** Annotated circular map of the *T. tuberosum* plastid assembly.

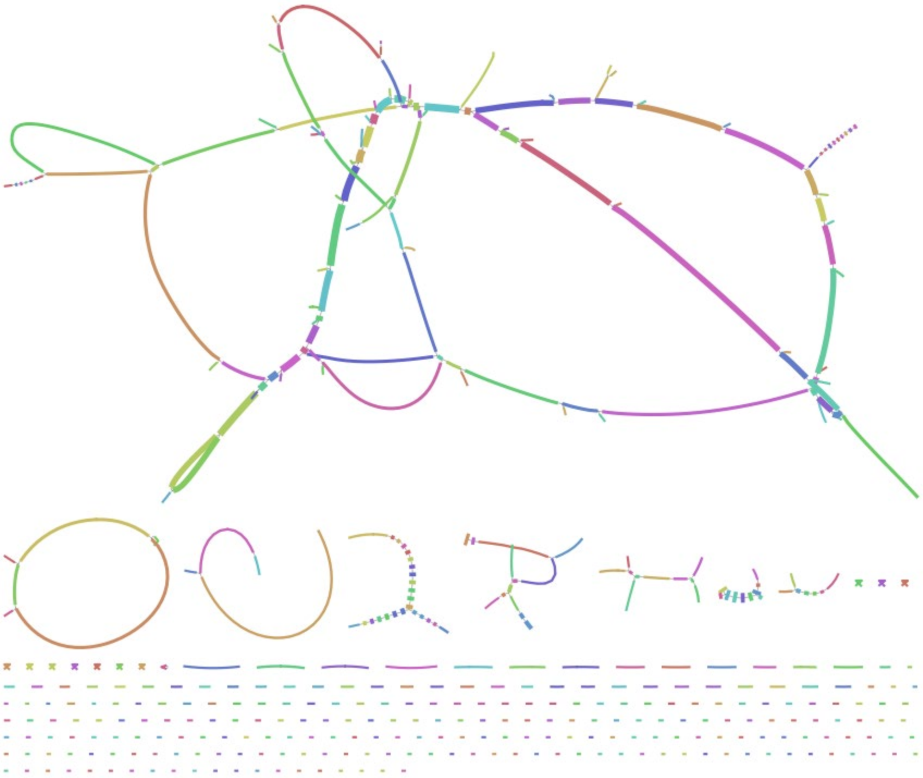

**Supplementary Figure 11:** Bandage plot of the *T. tuberosum* mitochondrial genome assembly illustrating assembly complexity. The plot represents one connected component of ~550 kb with complex repeat-mediated branching typical of plant mitochondrial genomes. Colors are random.

### Supplementary Tables

**Supplementary Table S1:** Statistics for the primary assembly of the *T. tuberosum* genome. Following removal of contaminants (n = 292), plastid-derived (n = 60), mitochondria-derived (n = 30) and repeat contigs (n = 2) from the raw primary assembly, BUSCO completeness confirms that no nuclear gene content was lost during filtering.

| | **Raw assembly** | **Filtered assembly** |
|---|---|---|
| Number of contigs | 2,189 | 1,805 |
| Total assembly size | 1.32 Gbp | 1.30 Gbp |
| Largest contig | 60 Mbp | 60 Mbp |
| N50 contig length | 32.2 Mbp | 32.2 Mbp |
| L50 contig count | 16 | 16 |
| N90 contig length | 1.1 Mbp | 1.3 Mbp |
| L90 contig count | 96 | 84 |
| GC content | 36.73 % | 36.70 % |
| BUSCO v5 embryophyta_odb10 | C:98.5% [S:21.7%, D:76.8%], F:0.7%, M:0.7%, n:1614, E:1.9% | C:98.5% [S:21.7%, D:76.8%], F:0.7%, M:0.7%, n:1614, E:1.9% |

**Supplementary Table S2:** Statistics for the pseudo-haplotype-resolved assembly of the *T. tuberosum* genome. Following removal of contaminants ($n_{hap1}$ = 301; $n_{hap2}$ = 18), plastid-derived ($n_{hap1}$ = 69; $n_{hap2}$ = 58), mitochondria-derived ($n_{hap1}$ = 67; $n_{hap2}$ = 36) and repeat contigs ($n_{hap1}$ = 6; $n_{hap2}$ = 13) from the raw pseudo-haplotype-resolved assembly, BUSCO completeness confirms that no nuclear gene content was lost during filtering.

| | **Raw assembly** | | **Filtered assembly** | |
|---|---|---|---|---|
| | **Hap1** | **Hap2** | **Hap1** | **Hap2** |
| **Number of contigs** | 2425 | 671 | 2022 | 580 |
| **Total assembly size** | 1.21 Gb | 0.98 Gb | 1.19 Gb | 0.96 Gb |
| **Largest contig** | 49.8 Mb | 50.2 Mb | 49.8 Mb | 50.2 Mb |
| **N50 contig length** | 18.4 Mb | 26.3 Mb | 18.4 Mb | 26.3 Mb |
| **L50 contig count** | 19 | 15 | 19 | 15 |
| **N90 contig length** | 0.33 Mb | 0.96 Mb | 0.51 Mb | 1.1 Mb |
| **L90 contig count** | 204 | 96 | 167 | 89 |
| **GC content** | 36.78 % | 36.17 % | 36.75 % | 36.19 % |
| **BUSCO v5 embryophyta_odb10 (n:1614)** | C:96.3% [S:28.4%, D:67.9%], F:1.0%, M:2.7%, E:1.7% | C:95.4% [S:39.0%, D:56.3%], F:0.9%, M:3.7%, E:2.3% | C:96.3% [S:28.4%, D:67.8%], F:1.0%, M:2.7%, E:1.7% | C:94.2% [S:37.9%, D:56.3%], F:0.8%, M:5.0%, E:2.4% |
| **BUSCO v5 embryophyta_odb10 (n:1614), combined** | C:98.8%[S:1.1%,D:97.7%], F:0.6%,M:0.7%,E:1.3% | | C:98.8%[S:1.1%,D:97.6%], F:0.6%,M:0.7%,E:1.3% | |

**Supplementary Table S3:** Taxonomic classification of 292 low-coverage contigs (< 10×) identified as contaminants by Kraken2 against the standard RefSeq database, and classification of 92 high-coverage contigs (> 200×) by BLASTn against assembled organelle genomes. Percentages are of the 384 total removed contigs.

| Taxon | Group | Category* | Contigs | % |
|---|---|---|---|---|
| *Xanthomonas albilineans* | Gammaproteobacteria | BC | 106 | 27.6% |
| *Lactiplantibacillus pentosus* | Bacilli | BC | 85 | 22.1% |
| *Lactiplantibacillus plantarum* | Bacilli | BC | 1 | 0.3% |
| *Prochlorococcus marinus* | Cyanobacteria | BC | 30 | 7.8% |
| *Limnospira indica* | Cyanobacteria | BC | 11 | 2.9% |
| *Acidovorax sp. BLS4* | Betaproteobacteria | BC | 19 | 4.9% |
| *Candidatus Liberibacter africanus* | Alphaproteobacteria | BC | 17 | 4.4% |
| *Wolbachia pipientis* | Alphaproteobacteria | BC | 2 | 0.5% |
| *Candidatus Hodgkinia cicadicola* | Alphaproteobacteria | BC | 2 | 0.5% |
| *Vibrio coralliilyticus* | Gammaproteobacteria | BC | 1 | 0.3% |
| *Pseudomonas sp. Z1-29* | Gammaproteobacteria | BC | 2 | 0.5% |
| *Spiribacter sp. 2438* | Gammaproteobacteria | BC | 1 | 0.3% |
| *Streptomyces sp. T12* | Actinomycetes | BC | 2 | 0.5% |
| *Chlamydiaceae (unresolved)* | Chlamydiia | BC | 1 | 0.3% |
| *Homo sapiens* | Mammalia | HC | 3 | 0.8% |
| Plastid-derived contigs | — | P | 60 | 15.6% |
| Mitochondria-derived contigs | — | M | 30 | 7.8% |
| Unresolved repeat/rDNA | — | R | 2 | 0.5% |

**BC: Bacterial contamination; HC: Human contamination; P: Plastid; M: Mitochondria; R: Repeat/rDNA*

**Supplementary Table S4:** Base-level accuracy and *k*-mer completeness of raw and filtered assemblies evaluated using Merqury with a Meryl database derived from HiFi reads.

| | Raw assembly | Filtered assembly |
|---|---|---|
| **Primary assembly** | | |
| Primary QV | 59.3076 | 60.4025 |
| Primary completeness [%] | 75.4533 | 75.4455 |
| **Pseudo-haplotype 1** | | |
| Hap1 QV | 59.1690 | 60.3696 |
| Hap1 completeness [%] | 70.5199 | 70.4458 |
| **Pseudo-haplotype 2** | | |
| Hap2 QV | 65.4469 | 65.2169 |
| Hap2 completeness [%] | 66.0210 | 67.2160 |
| **Combined pseudo-haplotypes** | | |
| Hap1+Hap2 QV | 60.9849 | 62.2738 |
| Hap1+Hap2 completeness [%] | 95.1334 | 94.9066 |

**Supplementary Table S5:** Cross-genotype validation statistics

| | |
|---|---|
| **Mapping** | |
| ONT reads mapped (%) | 99.91 |
| Primary alignments mapped (%) | 99.68 |
| **Contig coverage (all 1,805 contigs)** | |
| Well-covered (≥10×) | 982 (54.4%) |
| High-copy (>200×) | 27 (1.5%) |
| Partial coverage (5–10×) | 288 (16.0%) |
| Low / absent coverage (<5×) | 508 (28.2%) |
| Low-coverage total length | 23.5 Mb (1.8%) |
| **Repeat content (nuclear, excl. high-copy contigs)** | |
| Genome-wide repeat fraction (length-weighted) | 71.3% |
| Repeat density — low-coverage contigs | 998 bp/kb |
| Repeat density — well-covered contigs | 952 bp/kb |
| **Gene content** | |
| Total annotated gene models (ANNEVO) | 56,354 |
| mRNAGene density — low-coverage contigs | 164.3 models/Mb |
| Gene density — well-covered contigs | 128.2 models/Mb |
| Gene models on low-coverage contigs | 3,446 (6.1%) |

*Contig totals sum to 1,805: 982 well-covered + 27 high-copy + 288 partial + 508 low-coverage. Nuclear repeat and gene analyses exclude the 27 high-copy contigs (1,778 contigs, 1,270 Mb). Weighted mean coverage of well-covered nuclear contigs: 16.4×.*

**Supplementary Table S6:** Repeat landscape of the *T. tuberosum* genome.

| | Number of elements | Percentage of sequence |
|---|---|---|
| **Retroelements** | **613,664** | **36.85%** |
| LINEs: | 54,466 | 3.28% |
| RTE/Bov-B | 296 | 0% |
| L1/CIN4 | 54,170 | 3.27% |
| LTR elements: | 559,198 | 33.58% |
| Ty1/Copia | 234,974 | 12.87% |
| Gypsy/DIRS1 | 201,459 | 15.64% |
| **DNA transposons** | **49,786** | **1.91%** |
| Hobo-Activator | 4,185 | 0.22% |
| MULE-MuDR | 40,307 | 1.57% |
| Tourist/Harbinger | 1,902 | 0.07% |
| **Rolling-circles** | **3,048** | **0.13%** |
| **Unclassified:** | **680,448** | **18.67%** |
| **Small RNA:** | **31,591** | **13.2%** |
| **Simple repeats:** | **188,502** | **0.7%** |
| **Low complexity:** | **39,269** | **0.15%** |

**Supplementary Table S7:** Comparison of Helixer and ANNEVO ab initio annotations of the *T. tuberosum* reference genome. Statistics are derived from GAQET2 analysis.

| | Helixer | ANNEVO |
|---|---|---|
| **NCBI_TaxID** | 147035 | 147035 |
| **Gene_Models (N)** | 84,059 | 56,354 |
| **Transcript_Models (N)** | 84,059 | 56,354 |
| **CDS_Models (N)** | 84,059 | 56,354 |
| **Exons (N)** | 405,266 | 271,743 |
| **UTR5' (N)** | 84,059 | 0 |
| **UTR3' (N)** | 83,529 | 0 |
| **Both sides UTR' (N)** | 83,529 | 0 |
| **Overlapping_Gene_Models (N)** | 4,557 | 59 |
| **Single Exon Gene Models (N)** | 16,224 | 20,126 |
| **Single Exon Transcripts (N)** | 16,224 | 20,126 |
| **Total Gene Space (Mb)** | 273.8 | 196.1 |
| **Mean Gene Model Length (bp)** | 3,257 | 3,480 |
| **Mean CDS Model Length (bp)** | 1,105 | 1,379 |
| **Mean Exon Length (bp)** | 290 | 286 |
| **Mean Intron Length (bp)** | 480 | 550 |
| **Longest Gene Model Length (bp)** | 57,416 | 57,141 |
| **Longest CDS Model Length (bp)** | 16,221 | 16,227 |
| **Longest Intron Length (bp)** | 36,246 | 28,705 |
| **Shortest Gene Model Length (bp)** | 104 | 201 |
| **Shortest CDS Model Length (bp)** | 1 | 2 |
| **Shortest Intron Length (bp)** | 30 | 5 |
| **Models with early STOP** | 0 | 0 |
| **Models START missing** | 0 | 0 |
| **Models STOP missing** | 504 | 0 |
| **Models START & STOP missing** | 0 | 0 |
| **BUSCO embryophyta_odb10** | C:98.1% [S:23.2%, D:74.8%], F:1.2%, M:0.7%, n:1614 | C:98.3% [S:24.2%, D:74.2%], F:0.7%, M:0.9%, n:1614 |
| **BUSCO viridiplantae_odb10** | C:98.8% [S:24.0% ,D:74.8%], F:1.2%, M:0.0%, n:425 | C:98.8% [S:26.1%, D:72.7%], F:1.2%, M:0.0%, n:425 |
| **PSAURON SCORE** | 87.5 | 97.2 |
| **DETENGA_FPV** | T: 84,059; PcpM0: 56,694; PteM0: 134; PchM0: 8; PcpMte: 522; PteMte: 855; PchMte: 227 | T: 56,354; PcpM0: 49,618; PteM0: 41; PchM0: 3; PcpMte: 64; PteMte: 152; PchMte: 144 |
| **DETENGA_FP%** | T: 84,059; PcpM0: 67.45 ;PteM0: 0.16; PchM0: 0.01; PcpMte: 0.62; PteMte: 1.02; PchMte: 0.27 | T: 56,354; PcpM0: 88.05; PteM0: 0.07; PchM0: 0.01; PcpMte: 0.11; PteMte: 0.27; PchMte: 0.26 |
| **OMArk Consistency Results** | Cons:71.54% [P:10.31%; F:3.68%], Inco:8.50% [P:2.74%, F:1.03%], Cont:0.00%, Unkn:19.95% | Cons:90.45% [P:7.27%; F:2.32%], Inco:6.95% [P:1.54%, F:0.34%], Cont:0.00%, Unkn:2.60% |
| **OMArk Completeness Results** | malvids-HOGs: 11,704; S:34.57%, D:59.69% [U:59.49%, E:0.20%], M:5.74% | malvids-HOGs: 11,704; S:34.71%, D:59.25% [U:59.02%, E:0.23%], M:6.03% |
| **OMArk Species Composition** | rosids: 80.05% | rosids: 97.40% |
| **ProteinsWithTREMBLHits (%)** | 78.18 | 96.15 |
| **ProteinsWithSWISSPROTHits (%)** | 62.03 | 80.39 |
